# Cortical integration of tactile inputs distributed across timescales

**DOI:** 10.1101/2024.07.22.604577

**Authors:** Wenyu Wan, K. Richard Ridderinkhof, Arko Ghosh

## Abstract

Sensory experiences in the real world cut across timescales from milliseconds to seconds. Emerging evidence suggests that somatosensory processing is sensitive to the temporal structure of the stimuli in the sub-second scale, yet only a few select ranges within this scale have been studied. To process real-world information, the integration of tactile inputs must occur over a much broader temporal range. To address temporal integration across timescales, we recorded scalp EEG signals from somatosensory cortex in response to a train of tactile pulses presented to the fingertips with varying inter-stimulus intervals (ISI) spanning 100 to 10,000 ms. To capture a rich variety of influences of the temporal structure on the cortical signals, we used a multi-dimensional event-related potential where the stimulations are separated according to the next interval structure. We tracked cortical tactile processing through its early (<75 ms), intermediate (75 to 150 ms) and late stages (150 to 300 ms). We find that the early and late stages of cortical activity were similarly dominated by the preceding ISI; EEG signals were suppressed with ISIs < 500 ms and enhanced with longer ISIs, with this effect persisting even when ISIs were approximately 8 seconds. The intermediate stage of cortical activity was sensitive to both the previous and the penultimate ISIs. Our findings suggest that the specific somatosensory cortical networks integrate temporal structure across timescales to enable complex sensory experiences.

## Introduction

The peripheral nervous system encodes texture by using the timing of tactile inputs in the millisecond scale (Weber et al., 2013). In the central nervous system, the cortex is capable of separating sequences of tactile information according to their temporal pattern in the scale of milliseconds to seconds (Bale et al., 2017). This capacity may serve diverse sensory functions – from localising touch to learning behaviourally relevant patterns (Godde et al., 2000; Nguyen et al., 2016). In the real world, the temporal patterns cut across various time scales. For instance, the inter-touch interval distribution captured on the smartphone touchscreen is heavy tailed ranging from milliseconds to hours (Pfister and Ghosh, 2020). The somatosensory cortex may separate rich tactile sequences according to the recent history of tactile events – i.e., the previous and the penultimate intervals.

There is scattered evidence that the previous interval shapes somatosensory cortical processing, but the role of the penultimate interval is less clear. When the interval to the previous tactile stimulation is shortened, the somatosensory cortical response is attenuated Espenhahn et al. (2020). A similar pattern of results is also found for studies using electrical stimulation of peripheral nerves (Kisley and Cornwell, 2006; Spooner et al., 2020). The measurements are usually limited to few select intervals (such as continuous tactile stimulations separated by 150 and 1050 ms in Espenhahn et al. (2020) and paired tactile stimulations separated by 30, 60 and 150 ms in Wühle et al. (2011)). For very short (<150 ms) intervals, the suppression may be under the influence of cortical excitability, with higher attenuation corresponding to lower excitability (Lenz et al., 2012; Shagass and Schwartz, 1964). By using MEG and electrical stimulation, the signal attenuation has been localised to the primary and secondary somatosensory cortex (Hamada et al., 2002). The primary somatosensory cortex shows attenuation for intervals up to 100 ms, whereas the secondary somatosensory cortex shows attenuated signals for intervals up to 500 ms. A similar pattern of results is found for tactile stimulation, albeit the exploration only spanned intervals of 30 to 150 ms (Wühle et al., 2011). These results suggest that the tactile information lingers longer in the intermediate stage of cortical processing than in the early stage. The sustained information processing of preceding stimuli in the intermediate stage may allow both the previous and penultimate pulses to shape the somatosensory processing of the current pulse, as well as its modulation by the length of the respective inter-stimulus intervals, as long as the total sequence occurs within 500 ms. If and how the previous and penultimate tactile pulses at intervals spanning >500 ms shape the cortical processing is not clear.

The conventional analytical framework – involving only a few select intervals – is limited in its ability to address the role of rich temporal statistics in cortical processing. Furthermore, this framework can only capture a simple relationship between the temporal statistics and brain responses (e.g., the shorter the interval, the higher the cortical signal attenuation). By contrast, the framework of joint-interval distribution (JID) allows us to separately measure tactile experiences according to their next-interval dynamics (i.e., the duration of the previous interval relative to the duration of the penultimate interval). This framework has been previously used to capture the rich temporal statistics of spiking neurons (Eagan and Partridge, 1989; Fitzurka and Tam, 1999) and smartphone touchscreen interactions (Ceolini et al., 2022a; Ceolini and Ghosh, 2023; Ceolini et al., 2022b; Duckrow et al., 2021). According to this framework the stimulation triad can be separated according to the joint properties of the previous interval (say *k*) and the penultimate interval (say *k*-1), and then gathered in two-dimensional bins (b_n_ × b_n_ number of bins) spanning the JID. Here, an event-related potential (ERP) signal that reflects somatosensory (Espenhahn et al., 2020) can be estimated separately for each of the two-dimensional bins (henceforth called JID-ERP). This approach can capture more complex relationships than when using only few select intervals. For example, patterns associated with short consecutive intervals may be differently attenuated compared to the patterns associated with short followed by long intervals.

The different stages of sensory cortical processing may be distinctly influenced by these temporal statistics. This influence can be captured using the signal latencies of the ERPs derived from the separate two-dimensional bins of the JID. Therefore, this JID-ERP framework can in theory capture a rich variety of influences of the temporal statistics on brain activity. To reveal these typical dynamics, the high-dimensional feature space (b_n_ × b_n_ bins of the JID-ERP × T signal processing latencies of the ERP) can be reduced by using non-negative matrix factorization (NNMF) (Lee and Seung, 1999). NNMF is a dimensionality reduction technique that can help reveal hidden patterns in the data by decomposing a matrix into lower-rank matrices (not unlike principal components, but resulting in additive rather than orthogonal components). The reduction can yield low-dimensional and interpretable prototypical patterns (so-called meta patterns). This method of reduction has been used to reduce a range of complex datasets from identifying facial features in images to human behavioural patterns captured on the smartphone (Ceolini and Ghosh, 2023; Lee and Seung, 1999). Here, the method can reduce the JID-ERP into a few interpretable prototypical temporal structures (*meta-JIDs*) and their corresponding timing (*meta-ERPs*). The *meta-JIDs* represent how the cortical signals are modulated by preceding tactile pulses as a function of the penultimate and previous intervals, whereas the *meta-ERPs* represent how the patterns of modulations are distributed over the different signal processing stages.

In this study, we captured the EEG signals evoked by tactile pulse stimulations separated by a range of previous and penultimate intervals spanning 100 to 10,000 ms. The stimulations were presented at the fingertips. We used the JID framework yielding 5 × 5 two-dimensional bins. The ERPs spanned an epoch from –200 ms prior to the tactile pulse to 299 ms following the pulse. This yielded a combined JID-ERP feature space of 5 × 5 × 500 two-dimensional bins. We reduced the JID-ERP after stimuli onset (5 × 5 × 299 two-dimensional bins) into interpretable prototypical patterns by using NNMF. To preview, these *meta-JIDs* and *meta-ERPs* revealed that the intermediate stages of cortical processing play a role that stands in contrast to the early or late stages.

## Methods

### Participants

This study is part of a large data collection effort involving long-term smartphone behavioral logs and EEG in the laboratory. Participants were recruited from a pool accumulated through the agestudy.nl data collection platform. This platform leverages the Dutch brain registry (Zwan et al., 2021). Sixty-four healthy participants (age range: 20-81; 40 female) were enrolled. However, the self-reported health status of 4 participants altered leading to eliminations (one had tinnitus, one had stroke, one had arthrosis, and one had a psychiatric condition). All participants provided the informed consent and the study was approved by the Ethics Committee of Psychology, Leiden University (ERB-reference number: 2020-02-14-Ghosh, dr. A.-V2-2044).

### Tactile Stimulation

We used tactile stimulations while recording the EEG signals (see below). Continuous train of tactile stimuli were applied to participants’ three fingertips (right thumb, index finger, and little finger, in different blocks) by using miniature electromagnetic solenoid-type stimulator (Dancer Design, North Yorkshire, England). The stimulator was controlled by using the parallel port of a computer and MATLAB (MathWorks, Natick, USA). We delivered 1800 stimulations, with each stimulation lasting 10 ms-long and the stimulations were suprathreshold according to verbal reporting conducted while testing the attached stimulators. The stimulations were time-stamped by the EEG amplifier based on the cop (see below). The inter-stimulus interval varied from 100 to 10,000 ms. In order to cover any noise generated by tactile stimulators and make participants feel less bored during measurement, we played a documentary series (David Attenborough’s Africa) on a screen and delivered the audio stream via a neck speaker. Participants were encouraged to have a short break every ∼15 minutes and the stimulation session lasted about 1 h.

### EEG recordings and preprocessing

EEG data was collected by using 64 channels EEG caps (62 scalp electrodes, 2 ocular electrodes) with customized equidistant layout (EasyCap Gmbh, Wörthsee, Germany)(Haenzi et al., 2014). The EEG signals were gathered referenced to the vertex and amplified with BrainAmp amplifier (Brain Products GmbH, Gilching, Germany). The signals were recorded and digitalized at the sampling rate of 1KHz. The participants arrived for the measurement with washed hair and scalp, and we further degreased the skin at the contact sites by using alcohol swabs. We applied Supervisc gel (Easycap) to obtain an electrical contact between the skin and electrode, and targeted an impedance under 10 kOhm for each electrode.

The data was processed using the EEGLAB toolbox (Delorme and Makeig, 2004) on MATLAB 2022b (MATLAB, Mathworks, Natick). Any recorded electrode with impedance >10 kOhm was excluded. The data was band filtered between 0.1 and 75 Hz. Eye-blink related EEG signal artefacts were removed by using independent component analysis (Pontifex et al., 2017). Next, the data was further filtered between 1 and 45 Hz. An excluded channels were subsequently interpolated. The data was epoched using a temporal window of 500 ms: - 200 to 299 ms from the onset of the stimuli. All epochs with signals beyond the set ±80 μV threshold were excluded. The data was re-referenced to the average from scalp electrodes. Note, the baseline was corrected for each epoch by subtracting the signal mean from -200 to -50 ms. The electrode with the largest short-latency signals (latency of 25-75 ms) placed over the left somatosensory cortex and with longer latency signals (latency of 150-300 ms) located in the central areas were chosen for each participant.

We excluded 2 participants due to corrupted EEG data files and six participants were further removed based on a signal amplitude threshold (10^th^ percentile of the population distribution). This yielded 52 participants for the subsequent ERP analysis.

### JID-ERP

We adapted the joint-interval distribution (JID) approach to separate the tactile stimulations according to their next interval dynamics. We introduced this approach for the study of real-world touchscreen behavioral dynamics (Ceolini and Ghosh, 2023; Duckrow et al., 2021) .

Here, a previous interval (say *k*) is described in conjunction with the penultimate interval (say *k*-1). The tactile joint intervals (stimulation triads) were binned into 5 × 5 two-dimensional bins with edges (upper bound) of 0.5, 1, 2, 4, 8 s. Then, for each bin separately we estimated the tactile-related potential (ERP) by averaging the signals evoked by the last event of the triad. The threshold number of events necessary to estimate the ERP was set to 10. After these steps were conducted for a given individual, for some of the analysis we additionally pooled across the individuals by creating a grand-average of the binned ERPs. Note, to reveal whether a particular bin’s amplitude was higher or lower than other bins at a given latency of the ERP, the JID-ERP was normalised by using *z*-score at each of the latencies.

### Mass-univariate statistical analysis

To reveal which two-dimensional bins were consistently suppressed or facilitated, we used mass-univariate one-sample *t*-tests of the *z*-scored amplitudes at all the time points. Multiple comparisons were corrected by using false discovery rate (FDR, using fdr() from eeglab toolbox, Delorme and Makeig, 2004) applied across the entire JID-ERP (α = 0.05). To quantify the similarities of the JID-ERP at any given latency to the rest of the latencies, a cross-correlation analysis was employed based on *z*-transformed grand-averaged JID-ERP using the Spearman method. Multiple comparisons were corrected by using the FDR method (α = 0.05). The values along the diagonal of the cross-correlation matrix (ρ = 1, window of 25 ms from the diagonal) was eliminated from the FDR.

### Interpretable data-driven dimensionality reduction

In the mass-univariate analysis above, each two-dimensional bin of the JID-ERP is considered independently from the other bins. However, the patterns in the JID-ERP may simultaneously encompass various bins in a complex manner. To extract such patterns we leveraged non-negative matrix factorization (NNMF). This method was originally demonstrated in facial image analysis, revealing key facial features (Lee and Seung, 1999), and extended to the study of behavioral JID to discover prototypical behaviours (Ceolini and Ghosh, 2023). Here we applied a variant of the method, stable and reproducible NNMF (star NNMF) described here (Ceolini and Ghosh, 2023). In brief, at the level of each individual, we applied the NNMF on a matrix of JID-ERP, where the EEG signal magnitudes (absolute value) was *z*-scored at all ERP latencies. Because this method requires non-negative inputs, we subtracted minimum of the *z*-transformed matrix from the entire matrix. The dimensional reduction method yielded a lower-rank approximation in terms of prototypical patterns: how the signals were modulated across the two-dimensional bins, *meta-JIDs*, and the corresponding latencies where the pattern was present, *meta-ERPs*. In the search for the optimal cluster between 2 to 10 ranks, and evaluated the optimal cluster number by calculating the error between real matrix and reconstructed data. The stable reduction was established based on 100 repetitions. We further clustered similar *meta-ERPs* across the population by using *k*-means; optimal cluster size search between 3 to 7 clusters, 1000 repetitions, with *silhouette* method using *evalclusters*.*mat* as implemented in MATLAB. The NNMF was applied also applied to the grand-averaged JID-ERP matrix using the same steps as described above but the *k*-means clustering was not applicable in this case.

## Result

### Tactile evoked potentials and the distinct stages of cortical processing

This report focused on the high-dimensional JID-ERP approach, but we first described the simple event-related potential signal disregarding the next-interval dynamics to provide an overview of the signal components observed here. Note, throughout this section we focused on the findings from the stimulations presented at the index fingertip (with other stimulation sites described in the supplementary information). In response to tactile pulse stimulation at the index fingertip, the evoked responses showed short-latency signals at the electrodes placed over the somatosensory cortex, while longer-latency signals were observed over the central electrodes (Supplementary Movie 1 & 4). As expected, the short-latency signals from the electrodes located in somatosensory cortex showed components with peak latencies of ∼50 ms (positive peak), ∼75 ms (negative peak), ∼100 ms (positive peak), ∼150 ms (negative peak), and ∼200 ms (positive peak). For subsequent analysis using the JID-ERP approach, we focused on the electrode with the largest amplitude (latency of 25-75 ms) over the somatosensory cortex (Fig. 1B). In the rest of the report we use the terms *early, intermediate* and *late* stages of the sensory processing to refer to the latencies of 1-75, 75-150, and 150-300 ms, respectively. For the ERP stemming from the central electrode see Supplementary Fig. 1. Across both the somatosensory and central electrodes, similar patterns as for the index finger stimulation were also captured in response to stimulation of the thumb and index finger (Supplementary Fig. 2, 3, 4, 5).

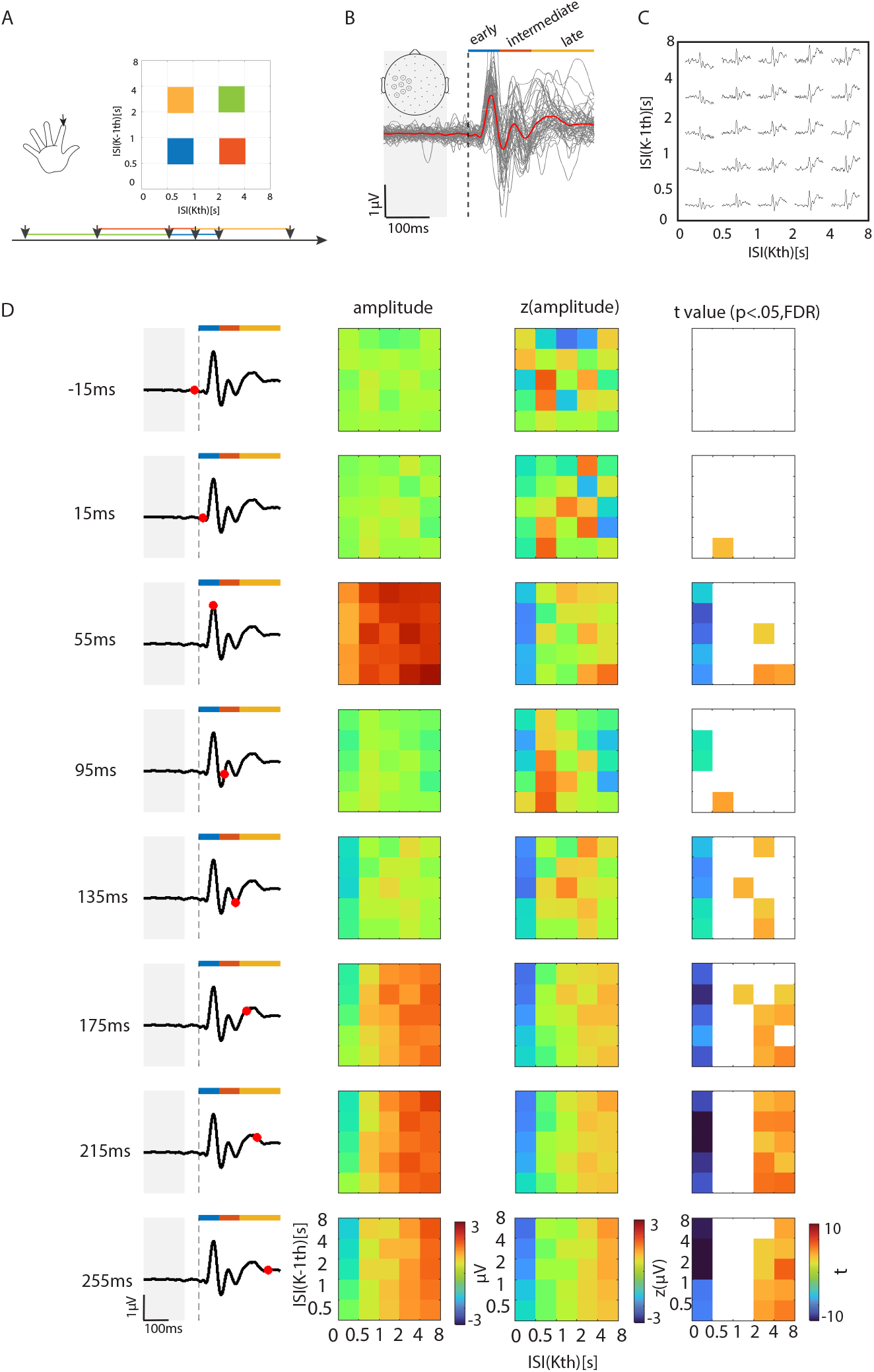

### Mass-univariate analysis reveals the influence of recent temporal statistics on tactile evoked potentials

To address how the tactile temporal statistics shapes the EEG signals, we separated the tactile ERPs according to the next-interval dynamics preceding the stimulation (JID-ERP). Here, a matrix of ERPs was separated in two-dimensional bins, binned based on the previous and the penultimate intervals (Fig. 1C). We analysed the differences in the ERP signal amplitudes across the JID for all the latencies of the ERP (JID-ERP). At each latency, the amplitudes were *z*-scored across the JID, and one-sample *t*-tests (*z*-scores vs. 0) revealed the two-dimensional bins with distinctly higher or lower activations (Fig. 1D). Note, in this approach the bins are considered independently of each other (mass-univariate statistics). In the pre-stimulation (but post ERP baseline) period, there was no consistent difference between the two-dimensional bins. The differences became evident at the early stages (∼50 ms post stimulation), such that the bins with short previous intervals <500 ms showed attenuated signals compared to the rest of the bins. The attenuation appeared irrespective of the penultimate interval. At this stage, there was evidence for signal enhancement for bins composed of short (<500 ms) penultimate intervals followed by long (>2,000 ms) intervals. The subsequent stage of processing, the intermediate stage, showed a more dynamic pattern of signal modulations which altered from one ERP peak to another. The late stage showed a pattern of modulations similar to the early stages – where the two-dimensional bins with <500 ms previous intervals showed attenuated signals. In this stage, there was enhancement of the signals with previous intervals >2,000 ms. A similar pattern of results was observed when the thumb or the little finger was stimulated (Supplementary Fig. 2 & 4, Supplementary Movie 2 & 3).

We also analysed the ERPs over the central electrode, consisting of a broad signal peaking at ∼ 200 ms. This entire signal was attenuated at the two-dimensional bins with previous interval <500 ms. This pattern was observed for all three stimulation locations (Supplementary Movie 4, 5 & 6, Supplementary Fig. 1, 3 & 5).

### Similarities in the pattern of JID-ERP modulation at the early and late stages of somatosensory processing

The JID-ERP revealed modulation of the EEG signals according to the two-dimensional bins based on the previous and penultimate intervals. As described above, the modulation varied according to the stage of processing for the somatosensory signals. We quantified the similarities of the JID-ERP at any given latency to the rest of the latencies by using correlations (Fig. 2A). For the index finger stimulations, this analysis revealed strong similarities between the JID-ERP patterns at the early and late stages, but the intermediate stage appeared largely unrelated to the other processing stages. There was a notable persistence of patterns within the different stages of processing – but not for the intermediate stages. The distinction of the intermediate stage was most prominent for the index finger, followed by the thumb and the little finger (Supplementary Fig. 2C & 4C). For the ERP over the central electrode, the JID-ERP patterns showed similar modulations across the intermediate and late stages (Supplementary Fig. 3C & 5C).

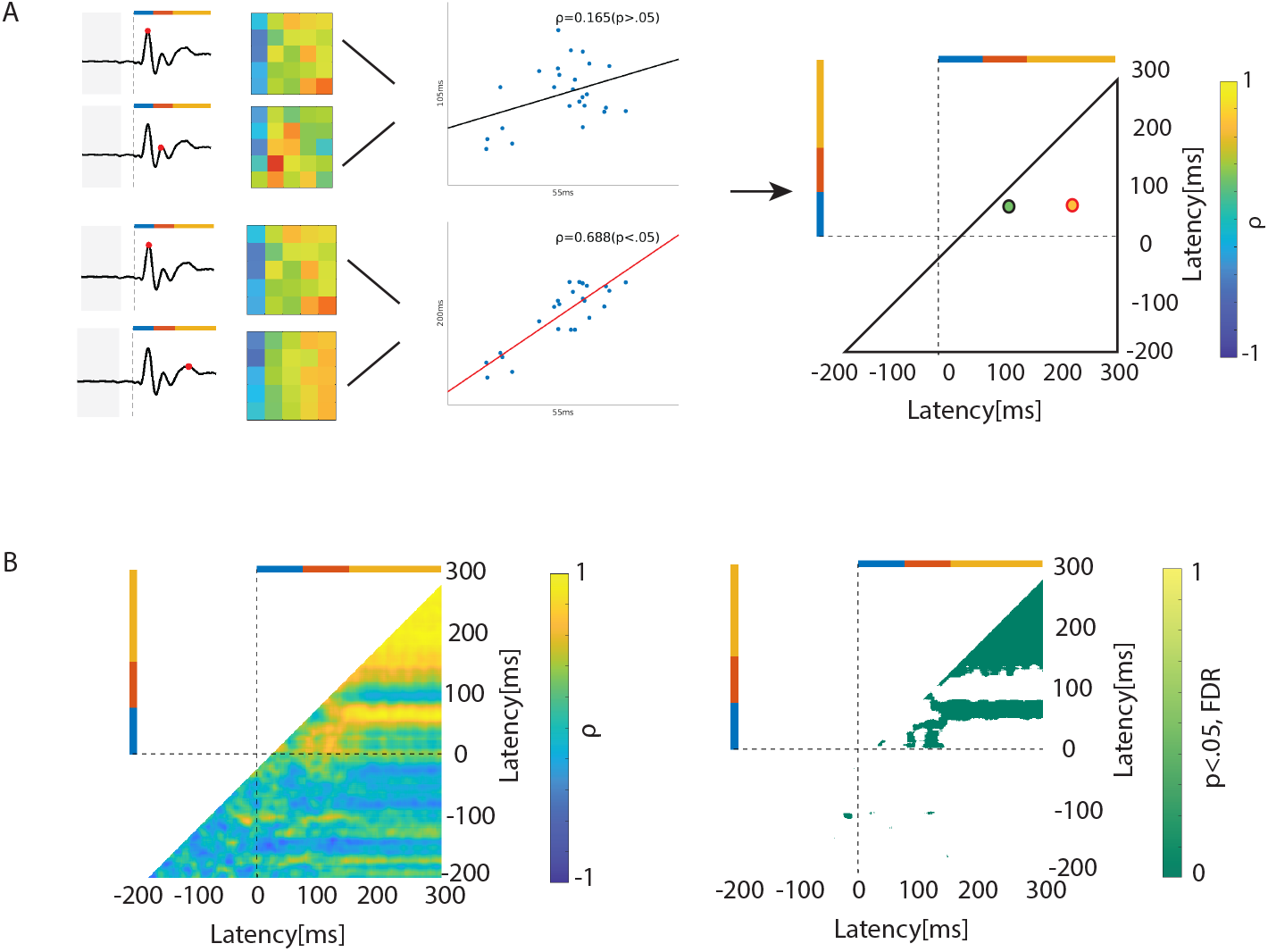

### Dimensionality reduction: Interval patterns relevant for somatosensory processing across the different stages of processing

By using simple analytical frameworks, i.e., mass-univariate statistics and correlation analysis, we established the dominant patterns underlying the JID-ERP, but these methods are not designed to extract time-variant complex patterns. Therefore, we next used data-driven dimensionality reduction to extract the key patterns of the rich JID-ERP. Essentially, the JID-ERP magnitudes (absolute value of the EEG signals, see Methods) spanning all the latencies were reduced in terms of a few *meta-JIDs*, i.e., prototypical interval-based modulations of the two-dimensional bins, and the corresponding *meta-ERPs*, i.e., the extent to which prototypical patterns were present at any given latency (Fig. 3A). We applied this analysis both on the grand averaged JID-ERP signal and on the individual JID-ERP traces, and also to the signals from the somatosensory and the central electrodes. Here we focus on the results from the index finger (for the thumb and little finger please see Supplementary Fig. 6 & 7).

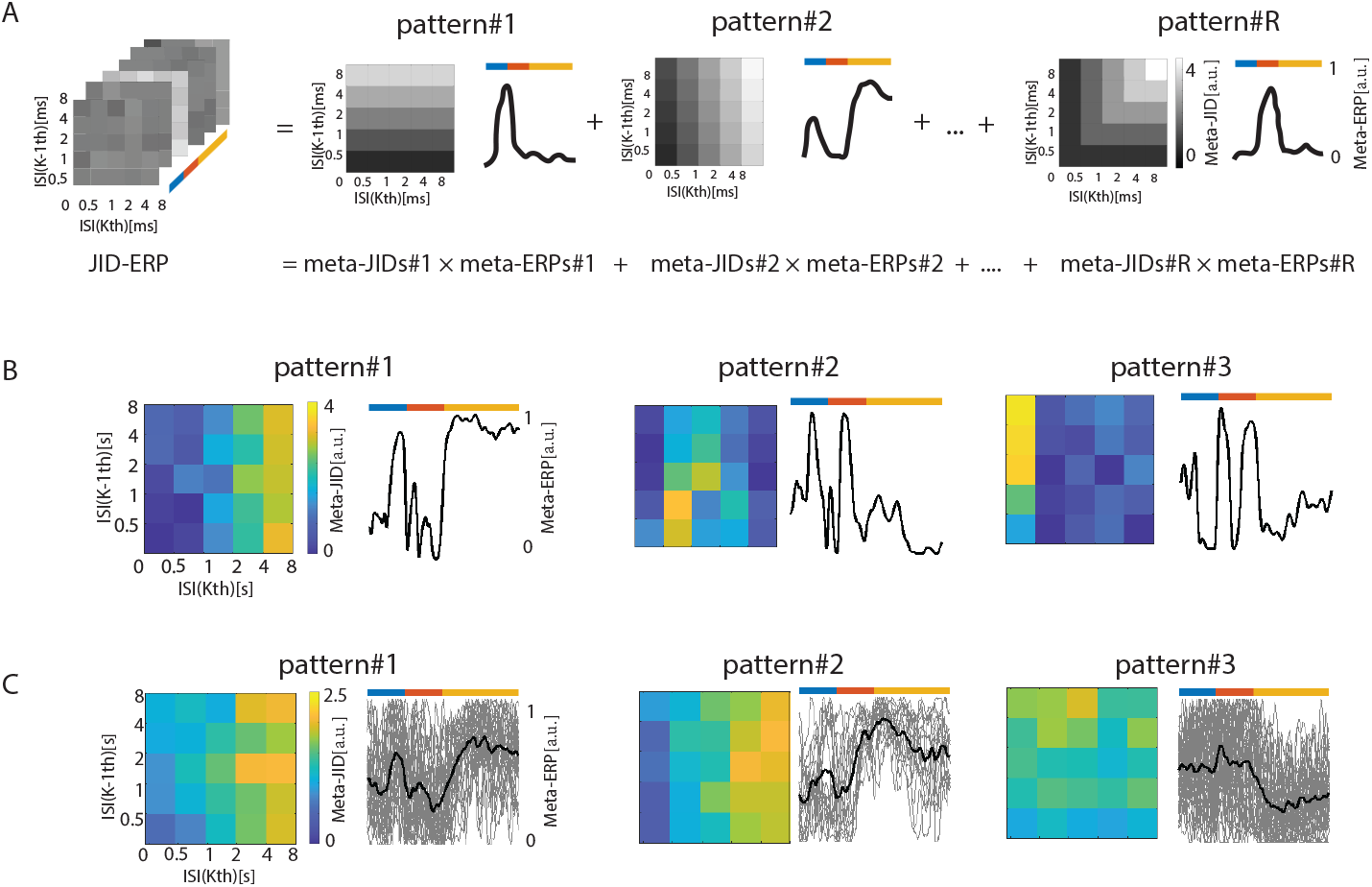

For the grand averaged signal from the somatosensory electrode, the dimensionality reduction revealed three prototypical patterns. First, we observed a prototypical pattern where short penultimate followed by short previous intervals were attenuated and the attenuation diminished for longer previous intervals. According to the corresponding *meta-ERPs* this pattern was present in the early and late stages of somatosensory processing. Second, we observed a more complex prototypical pattern where penultimate intervals <1,000 ms and previous intervals between 500 and 1,000 ms showed the larger amplitudes. This pattern was present in both the early and intermediate stages of processing. Third, we observed a pattern where previous intervals <500 ms showed larger amplitudes than longer previous intervals, and within this range, the longer penultimate intervals showed larger amplitudes. This pattern was largely constrained to the period of transition to and from the intermediate stage.

We performed a similar analysis at the level of each individual and then clustered the results based on the *meta-ERPs*. The corresponding *meta-JIDs* were averaged across the clusters. This approach revealed three prototypical patterns. First, we observed a pattern similar to the first prototypical pattern identified using the grand average signal. Here, the signals were attenuated for short consecutive intervals and enhanced for the temporally distant consecutive intervals. This pattern was present in the early and the late stages of somatosensory processing. Second, we observed a pattern similar to the first pattern but with a sharper attenuation for previous intervals <500ms and with strong presence in the late stages of the processing. Third, there were *meta-ERPs* which indicated peculiar modulation at the intermediate stages but the averaged *meta-JIDs* revealed only a marginal enhancement for long penultimate intervals followed by short previous intervals – the marginally central tendency is indicative of inter-individual variability at this stage.

The grand averaged signal at the central electrode revealed two prototypical patterns. The first was present only in the early stages of somatosensory processing. This pattern appeared complex with a tendency for enhanced processing of long penultimate intervals followed by short intervals (Supplementary Fig. 7A). The second was present in the intermediate and late stages. This pattern was characterised by an attenuation of stimulations with short previous intervals. The individual-participant signals revealed an additional complex prototypical pattern which was specific to the intermediate stage (Supplementary Fig. 7A).

## Discussion

We analysed cortical signals in response to a train of tactile inputs on the hand, with the interval between stimulations spanning ∼100 to 10,000 ms. Signal processing in the somatosensory cortex was shaped by both the penultimate and previous tactile pulses, and this influence was modulated by the respective inter-stimulus intervals spanning up to 8,000 ms. The intervals modulated the cortical processing by using rules which varied according to the stages of processing. By using a series of analytical approaches, we showed that the early and the late stages of cortical processing were similarly influenced by the interval patterns. In both of these stages, the signals were attenuated when the previous intervals were shorter than 500 ms. The influence of the temporal statistics at the intermediate stages was modulated by a more complex combination of the penultimate and previous intervals. Our findings indicate that the ability to separate tactile information according to temporal structures in the scales of milliseconds to seconds is distributed across different cortical networks.

One of the prominent patterns we observed included a sharp attenuation of the early cortical signals for previous intervals <500 ms. This is consistent with prior studies (Hamada et al., 2002; Zhu et al., 2007). For instance, inter-touch intervals separated by 2,000 ms resulted in larger amplitude in early stages of the ERP (<75 ms) compared to intervals separated by <500 ms; the late-stage ERPs (75-150 ms) were virtually abolished by the shorter intervals (Zhu et al., 2007). One possibility is that this attenuation reflects a form of inhibition induced when processing the first input (not unlike prepulse inhibition of startle responses; see, e.g., Braff et al., 2001). In our observations, the penultimate intervals played only a marginal role in further attenuating signals; the putative inhibitory processes appeared short-lived (i.e., lasts <500 ms). The data driven dimensionality reduction revealed an enhancement of the signal amplitudes at the offset of the early processing stage (latency of ∼75 ms). This may be explained by a rebound of activation, as often seen post-inhibition (Qi et al., 2016). Finally, there may be a concern that the attenuation observed does not reflect altered neural activity but rather how the EEG signal evoked by the second stimuli summates with the previous stimuli. However, this is unlikely as we saw no signal changes in the segment that appeared after the baseline period but prior to the second stimuli, and there was temporal jitter in the intervals within each bin.

Another pattern of modulation consisted of amplitude gradient across the ‘diagonal’ of the joint-interval distribution: with stronger signal attenuation for consecutive short-short intervals than for consecutive long-long intervals. The gradient spanned ∼100 to 8,000 ms. We speculate that this pattern reflects mechanisms that help sustain neural representations towards temporal integration of event series, such as short-term memory or perceptual priming (Contreras et al., 2013; Han et al., 2008). Animal studies suggest that this kind of reverberation of brain activity for past events can last up to a few minutes (Contreras et al., 2013). Based on the ERP signal latencies, we suggest that the activity in the early stages is echoed or recurred in the late stages resulting in the similarity of prototypical pattern between those two stages (Auksztulewicz et al., 2012; de Lafuente and Romo, 2006). Alternatively, some top-down modulation at the sensory cortex (Ruff, 2013) may exist in the late-stage processing to get ready for the arriving stimuli.

Our results also indicated a function of the intermediary stages which is distinct from the early and late stages of somatosensory processing. According to the correlational analysis, the JID-ERP was distinct at these stages as opposed to the early and late stages. Furthermore, the dimensionality reduction also revealed complex prototypical patterns largely unique to the intermediary stages. Our results are in line with previous studies indicating that complex temporal integration follows early somatosensory cortical processing (Rossi-Pool et al., 2021). However, exactly which mechanisms underscore these remains unclear. It is possible that stimulation induced inhibition, sustained neural representation, and links to extra-somatosensory processes (such as for motor control) all contribute to the distinct processing at the intermediary stages (Forss and Jousmaki, 1998; Haggard and Whitford, 2004).

In this study we separated the ERP signals according the dynamics of the tactile stimulations. The patterns captured in the so-called JID-ERP suggest that tactile processing is shaped according to the temporal context. This parallels the emerging view from both human and animal studies that cortical sensory processing is state-dependent (Parajuli et al., 2023; Russell et al., 2024; Taga et al., 2018; Zagha et al., 2013).

Our study provides a new perspective on studying how brain states interact with continuous sensory stimuli from outside. Our findings lead to the possibility that early stages of cortical processing may leverage the timing of previous and penultimate inputs to recognise the current state. Such a process may work in tandem with other feed-forward processes in early sensory processing to prioritise select input features (Westerberg et al., 2023). The sensory processing revealed here was largely consistent across the different fingers – the thumb, the index and little finger – indicating that the discovered state-dependent processing may generalise to the rest of the hand. The JID-ERP approach introduced here may be extended to other body parts or even other sensory modalities to reveal the shared mechanisms underlying state dependent cortical sensory processing.

## Data, Materials, and Software Availability

Pseudo-anonymized MATLAB.mat file containing processed JID-ERP data is made available on dataverse.nl upon publication; The custom-written scripts used towards this report are shared on: https://github.com/CODELABCODELIB/JID_ERP_Tactile_2024.

## Acknowledgements

This research was funded by Velux Stiftung (No.1283, awarded to A.G. with K.R.R as co-applicant). We thank Ruchella Kock for her help in programming. We thank Beste Yavuz, Lysanne Groenewegen, Barbora Michalidesová, Lorenzo Van Hoorde, and etc. for assistance in EEG data collection. We thank all participants for contributing with their time and effort.

## Author contributions

A.G. conceived the study. A.G. and W.W. designed the study. A.G. programmed the experiments and performed the EEG pre-processing. W.W acquired and analysed the data with the aid of A.G.. W.W drafted with the aid of A.G.. K.R.R, A.G., W.W. edited the manuscript.

